# Molecular architecture of a membrane-spanning hormone acyltransferase required for metabolic regulation

**DOI:** 10.1101/556233

**Authors:** Maria B. Campaña, Flaviyan Jerome Irudayanathan, Tasha R. Davis, Kayleigh R. McGovern-Gooch, Rosemary Loftus, Mohammad Ashkar, Najae Escoffery, Melissa Navarro, Michelle A. Sieburg, Shikha Nangia, James L. Hougland

**Author notes:** These authors contributed equally to the work (Maria B. Campaña, Flaviyan Jerome Irudayanathan). To whom correspondence should be addressed: Shikha Nangia, James L. Hougland.

## Abstract

Integral membrane proteins represent a large and essential portion of the proteome that often prove challenging for structural studies. We demonstrate a synergistic approach to structurally model topologically complex integral membrane proteins by combining co-evolutionary constraints and computational modeling with biochemical validation. We report the first structural model of a eukaryotic membrane-bound O-acyltransferase (MBOAT), ghrelin *O-*acyltransferase (GOAT), which modifies the metabolism-regulating hormone ghrelin. Our structure suggests an unanticipated strategy for trans-membrane protein acylation, with catalysis occurring in an internal channel as GOAT acts as an “enzyme inside a pore”. Our structure opens the door to structure-guided inhibitor design targeting GOAT and other MBOAT family members while validating the power of our approach to generate predictive structural models for other experimentally challenging integral membrane proteins.

## Introduction

Integral membrane proteins represent a large and essential portion of the proteome, including a growing number of enzymes, receptors, and transporters that serve as desirable drug targets.(*1*, *2*) However, these proteins often prove recalcitrant to purification and structural analysis due to their hydrophobic nature and reliance on interactions with lipid bilayers for both stability and activity.(*3*, *4*) We report here a synergistic approach to develop a structural model of topologically complex integral membrane proteins by combining co-evolutionary contact constraints and computational modeling with biochemical validation. Building upon only the primary sequence of the protein of interest and a biochemical assay for protein function, our approach provides an accessible and efficient route to build structural models of intractable membrane protein targets.

We demonstrate this approach by developing a structural model for ghrelin *O*-acyltransferase (GOAT), a member of the membrane bound *O*-acyltransferase (MBOAT) enzyme family responsible for octanoylation of the peptide hormone ghrelin (Fig. 1).(*5*, *6*) One of three protein-modifying MBOAT family members alongside Hedgehog acyltransferase (Hhat) and Porcupine (Porcn),(*7*–*9*) GOAT plays a central role in regulating energy homeostasis and metabolism through octanoylated ghrelin-dependent signaling pathways.(*10*) While the unique chemistry and biology of ghrelin and GOAT has inspired continued interest in targeting this system for therapeutic benefit, the inability to purify active GOAT and determine its structure has hampered progress towards this goal.(*11*–*13*) In this work, we report the first structural model for a eukaryotic MBOAT family member. Our human GOAT (hGOAT) structure is highly consistent with a recently reported crystal structure for the D-alanyl transferase DltB, a bacterial MBOAT homolog.(*14*) Our structure suggests a novel strategy for solving the topological challenge of trans-membrane protein acylation, with catalysis occurring in an internal channel within hGOAT without the octanoyl-CoA donor being transported into the endoplasmic reticulum (ER) lumen. The availability of this structure of hGOAT, a therapeutically interesting enzyme, opens the door to structure-guided inhibitor design targeting GOAT and other MBOAT family members while validating the power of our approach for creating molecular models for other experimentally challenging integral membrane proteins.

**Figure 1.**
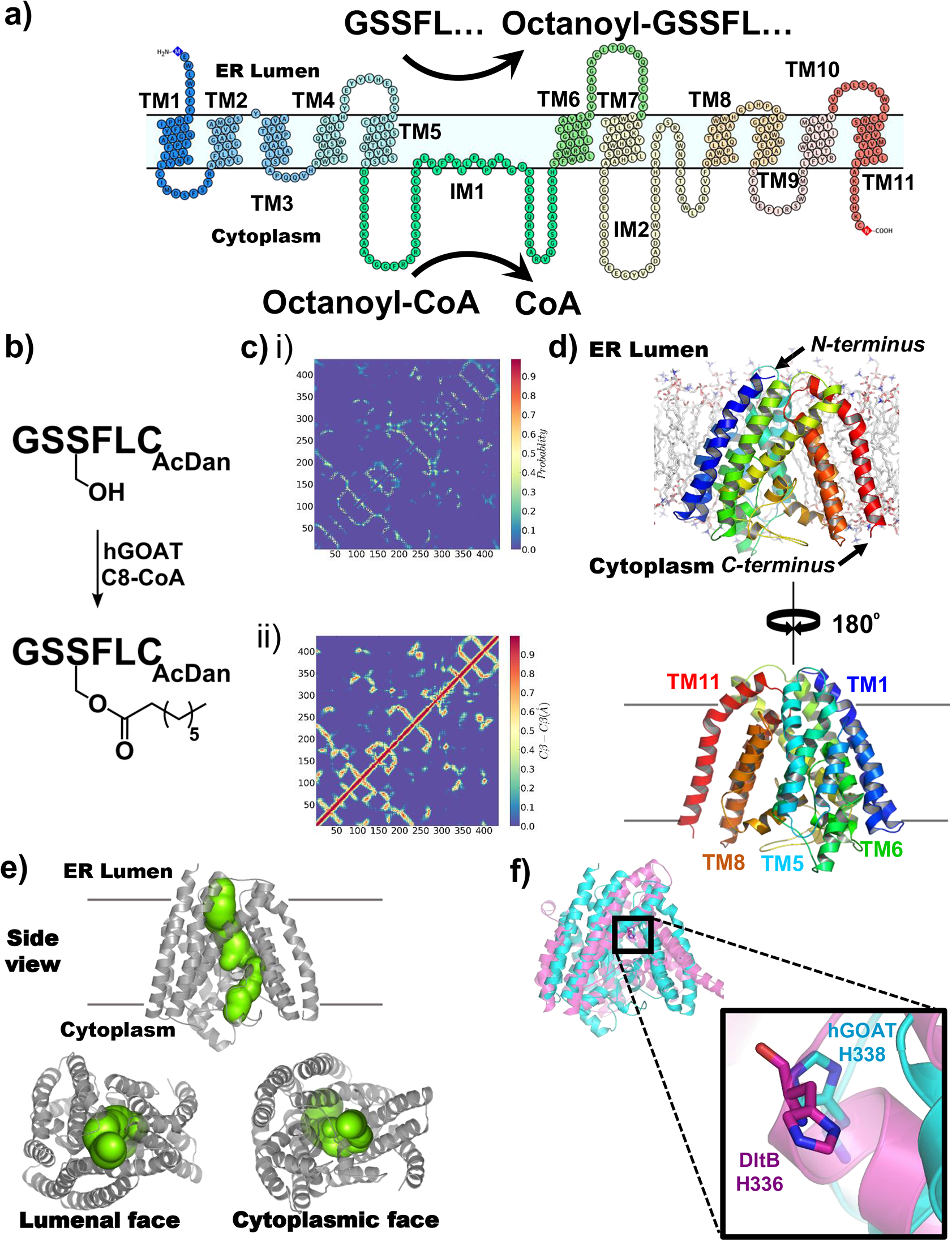
Structural model of hGOAT generated by computational methods. a) Schematic of ghrelin octanoylation by hGOAT showing the predicted transmembrane topology of hGOAT containing eleven transmembrane helix domains (TM1-11), two intramembrane domains (IM1-2), and loop regions generated using Protter.(*31*) b) Octanoylation of a ghrelin-mimetic fluorescent peptide by recombinant hGOAT. c) Contact maps for hGOAT showing the probability for a co-evolutionary contact from RaptorX analysis (i) and amino acid contacts in the final optimized hGOAT structure (ii). d) Structure of hGOAT in an ER-mimetic lipid membrane, correlated to color-coded membrane topology in part a). e) Illustration of internal channel within hGOAT (green) transiting from the ER lumen to the cytoplasm, with the channel determined by CAVER 3.0 plugin in PyMOL.(*32*) f) Structural overlay of hGOAT and DltB showing the absolutely conserved histidine residues (hGOAT H338, teal; DltB H336, purple, PDB ID 6BUG:C) within the cores of these acyltransferases.

## Development of a computational model for human GOAT structure

In generating our hGOAT structural model, we utilized state-of-art co-evolutionary contact predictions with computational protein folding and structure optimization methods (Supp. Fig. 1).(*15*, *16*) Using metagenomics protein databases, we generated a multiple-sequence alignment to predict residues that are potentially in contact (defined as C_β_-C_β_ < 8 Å) with each other in the folded structure (Supp. File S1).(*15*, *17*) This set of contacts (Supp. File S2), represented by the contact map (Fig. 1c and Supp. Fig. 2), guided our hGOAT structural modeling.(*17*–*19*) Experimental information on GOAT membrane topology and co-evolutionary contact constraints were iteratively combined in protein folding simulations to generate a manifold of ∼30,000 potential hGOAT structures.(*11*, *18*, *20*) The generated structures were clustered, and the lowest energy structures that satisfied the contact map were isolated.(*20*) Representative structures from the top five clusters were then subjected to further structural refinement to yield the optimal hGOAT model.(*21*) The optimal model was embedded in a lipid membrane and subjected to structural relaxation in explicit solvent using all-atom molecular dynamics simulations.(*22*–*24*) This simulation used an ER-mimetic lipid bilayer to ensure optimization of hydrophobic protein-lipid interactions.(*25*)

## Analysis of the predicted structure of human GOAT

Our computationally-derived structure for hGOAT is consistent with the previously reported topological model of the mouse GOAT ortholog,(*11*) containing a total of eleven transmembrane helices with slightly altered helix boundaries (Fig. 1a). The enzyme forms an ellipsoidal cone composed of transmembrane helices, with the narrow end facing the ER lumen (Fig. 1d). The exposed ends of five transmembrane helices (TM1, TM4, TM5, TM7, and TM11) converge to form a pore through which the interior of hGOAT is connected to the ER lumen. At the cytoplasmic membrane interface, the predicted cytoplasmic loops fold up to form a core region bounded by the lipid-contacting perimeter helices. As a result, there is minimal cytoplasmic exposure of hGOAT residues beyond the plane of the membrane, suggesting our description of hGOAT as an “enzyme inside a pore”.

The hGOAT structure contains a contiguous internal channel that transits from the ER to the cytoplasm through the enzyme core (Fig. 1e). The channel is bent within hGOAT, with the restriction formed by the C-terminal end of helix TM8 and N-terminal end of TM9. This positions an absolutely conserved histidine residue (H338) at the end of TM8 directly contacting the channel,(*7*) consistent with proposals for this histidine to serve as a general base for catalyzing ghrelin acylation. The H338 residue closely matches the location of the analogous histidine residue (H336) in the DltB structure (Fig. 1e).(*5*, *14*) Further comparison of the hGOAT model and the DltB structure (released during the preparation of this manuscript) reveals remarkable similarities in overall topology and structure, with an TM-Score of 0.6 and RMSD of 2.23 Å for aligned residues between the structural models for these distantly related MBOAT family members (12.3% sequence identity, 26.8% sequence similarity, E-Value 2.7×10^−8^ and bit score 48.7; Supp. Fig. S3 and Supp. Table S1).(*26*)

## Analysis of hGOAT structural model by alanine mutagenesis

To biochemically validate this computational hGOAT structural model, we mutated approximately 10% of the residues within hGOAT to alanine and determined the impact of these mutations on hGOAT octanoylation activity in a peptide-based acylation assay (Fig. 2).(*27*, *28*) These 42 alanine mutations were spread across a range of amino acids and degrees of conservation, with the majority of sites chosen conserved at >75% amongst GOAT orthologs (Supp. File S3). Approximately half of the mutation sites were selected based on the residue side chain contacting the internal void in our model, as we propose this channel within hGOAT will likely contain the substrate binding sites and catalytic residues.

**Figure 2.**
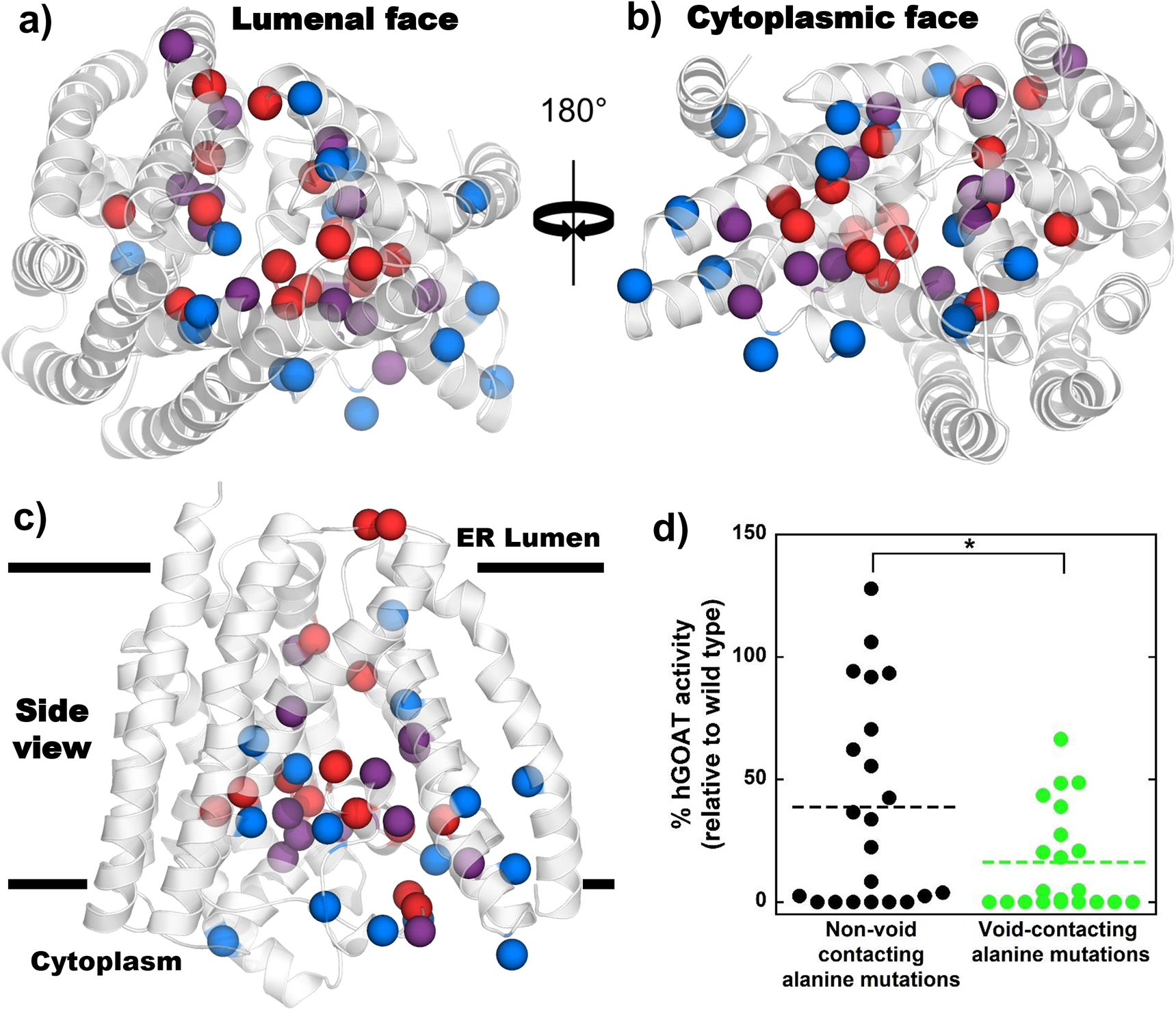
Mutagenesis studies support the location and functional importance of the hGOAT internal channel. a-c) Alanine mutations are mapped onto the hGOAT structure, with each sphere denoting the alpha carbon of the mutated residue; spheres are colored as follows: Blue, alanine variants with octanoylation activity within 3-fold of WT hGOAT; purple, alanine variants with impaired octanoylation activity (>3-fold loss compared to WT hGOAT); red, inactive alanine variants. a) View from the lumenal face; b) view from the cytoplasmic face; c) side view in the plane of the ER membrane. d) Octanoylation activity of hGOAT alanine variants for non-void contacting (black) and void-contacting mutations (green), with dotted lines denoting the average acylation activity for each group; *, p < 0.03.

Amongst this pool of alanine mutants, we observed a range of activities from near/above wild type to complete loss of detectable ghrelin octanoylation activity (Supp. Table S2 and Supp. Fig. S6). When mapped onto the hGOAT structural model, mutations leading to a marked decline (>3-fold, purple) or loss of enzyme activity (red) appear clustered within the core of hGOAT (Fig. 2). For quantitative analysis of the impact of these mutations, we determined whether alanine mutagenesis of residues contacting the internal void is more likely to yield reduced enzyme activity compared to non-void contacting mutations. Within the pool of mutations, the void-contacting alanine mutations were significantly more likely to result in loss of enzyme activity (p<0.03, Fig. 2d). This mutation activity mapping defines a functionally essential core within hGOAT consistent with our “enzyme inside a pore” model and expands the number of residues within hGOAT known to be required for enzyme activity.(*5*, *6*, *11*)

## Prediction and biochemical validation of the octanoyl-CoA binding site within hGOAT

We expect the octanoyl-CoA acyl donor to enter the hGOAT active site through interaction with the cytoplasmic face of the enzyme, based on the availability of acyl-CoAs within the cell. When docked into our hGOAT model, octanoyl-CoA binds to hGOAT through interactions of both its coenzyme A and octanoyl chain regions with residues in TM6, the TM7-TM8 connecting loop, TM8, and TM9 (Fig. 3). In this docked complex, the coenzyme A portion forms both polar and nonpolar interactions with multiple hGOAT residues while remaining exposed to the cytoplasm when bound to hGOAT (Fig. 3a-b). The phosphoadenosine group binds into a discrete pocket while the phosphopantetheine chain is in contact with multiple polar amino acid side chains (Fig. 3c-e). Amongst these CoA-contacting amino acids, all alanine mutations examined except one lead to a loss of hGOAT activity.

**Figure 3.**
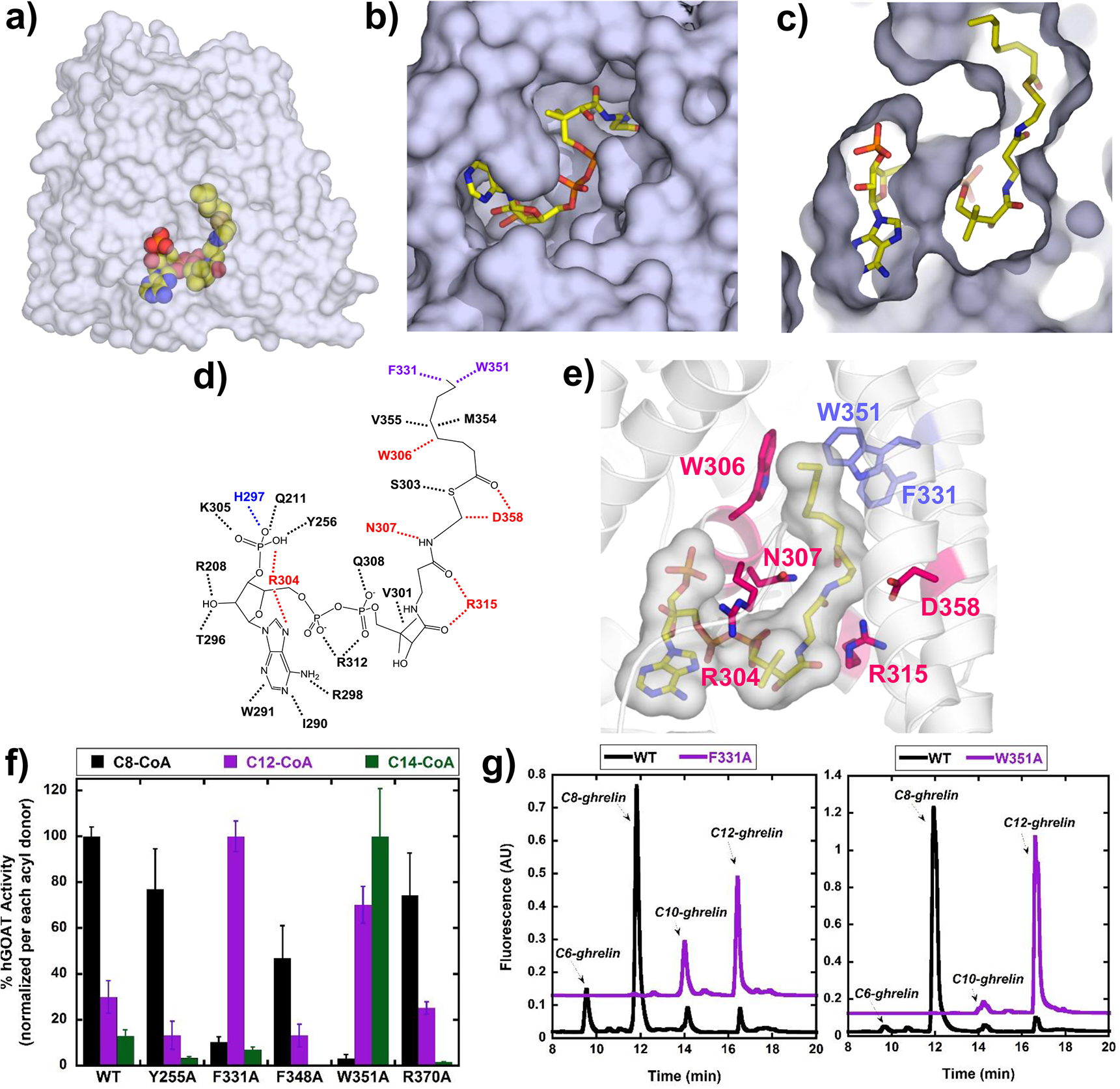
The acyl donor binding site within hGOAT. a) Structure of octanoyl-CoA bound within hGOAT from a side view in the plane of the ER membrane. b) View from the cytoplasmic face of hGOAT showing the solvent-exposed portions of the coenzyme A component of octanoyl-CoA. c) Cutaway view showing the acyl chain binding pocket within hGOAT, bent sharply upward from the coenzyme A binding regions on the cytoplasmic face of hGOAT. d) - e) Interactions between the octanoyl-CoA acyl donor and hGOAT residues; hGOAT residues shown in purple reduce acylation activity under standard reaction conditions when mutated to alanine, and residues shown in red abolish acylation activity upon alanine mutation. f) Acylation activity of WT hGOAT and selected hGOAT alanine variants using octanoyl-, lauryl-, or myristoyl-CoA as the sole acyl donor. Activities are normalized to the most reactive hGOAT variant with each acyl donor, and error bars reflect the standard deviation from a minimum of three independent trials. g) Acyl donor competition demonstrates altered selectivity to a longer acyl donor for F331A and W351A hGOAT variants, consistent with the predicted interaction of these amino acid side chains with the distal end of the octanoyl acyl chain.

In the docked hGOAT:octanoyl-CoA structure, the acyl chain of octanoyl-CoA makes a sharp turn and penetrates upward into the interior of hGOAT following a channel that terminates at W351 (Fig. 3c-e). Given the unique preference of hGOAT for an octanoyl acyl donor,(*6*, *13*, *29*) we examined alanine mutagenesis of predicted contacts within this acyl binding pocket to determine the impact of these mutations on hGOAT acyl donor selectivity. As alanine mutagenesis would provide additional space within the acyl binding site, we determined the ability of hGOAT alanine variants to accept twelve carbon (lauryl-CoA) and fourteen carbon (myristoyl-CoA) acyl donors in place of octanoyl CoA. The wild type enzyme and the majority of hGOAT alanine variants exhibit the expected preference for an eight carbon acyl donor, but alanine mutagenesis of W351 and F331 results in loss of appreciable reactivity with octanoyl-CoA and engenders new activity with the longer acyl donors (Fig. 3f and Supp Fig. S7). The F331A variant gains activity with the C12 donor while W351A hGOAT can acylate a ghrelin-derived peptide with both C12 and C14 acyl chains. This altered selectivity supports the modeled positions of W351 and F331 forming the end of the acyl binding pocket. The altered preference for longer acyl donors by the F331A and W351A variants is also observed in a direct competition assay where hGOAT variants are provided acyl donors ranging from six to twelve carbons. (Fig. 3g) This altered selectivity is not observed for any other alanine variants with detectable ghrelin acylation activity (Supp. Fig S8). Acyl donor reengineering upon targeted alanine mutagenesis localizes F331 and W351 to the distal end of the acyl donor binding site within hGOAT and provides further biochemical validation of our hGOAT structural model.

## Discussion and Conclusions

While structural studies play a central role in increasing our understanding of protein function, the challenge of structural modeling is particularly acute for integral membrane proteins given the limited availability of these proteins within structural databases.(*30*) In this work, we demonstrate the development and validation of a structural model for an integral membrane protein that leverages bioinformatics constraints from coevolutionary contact analysis and model evaluation by biochemical analysis while circumventing the requirement of protein purification.

Our model provides indispensable and novel insights into several long-standing questions regarding the mechanism for MBOAT-catalyzed transmembrane protein acylation. The topological separation of two essential conserved residues, H338 and N307, is explained by these two residues playing roles in distinct aspects of GOAT activity. The location of H338 within the central channel of GOAT, identical to the position observed for the analogous histidine in DltB,(*14*) is consistent with this residue acting as a general base to activate the ghrelin serine hydroxyl side chain for octanoyl transfer. In contrast, our hGOAT:octanoyl-CoA model implicates N307 in the binding site for the acyl donor. Based on our models of hGOAT and the hGOAT:octanoyl-CoA complex, we propose that hGOAT catalyzes transmembrane acylation of ghrelin by binding both substrates within the hGOAT internal channel and “handing off” the octanoyl group from CoA to ghrelin within this channel (Fig. 4). While many aspects of this proposed pathway such as the ghrelin binding site and location of catalytic residues remain to be functionally validated by ongoing studies, the established ability of our hGOAT model to efficiently guide biochemical studies demonstrates a novel approach to advance investigations of similar membrane proteins that are intractable to current structural approaches.

**Figure 4.**
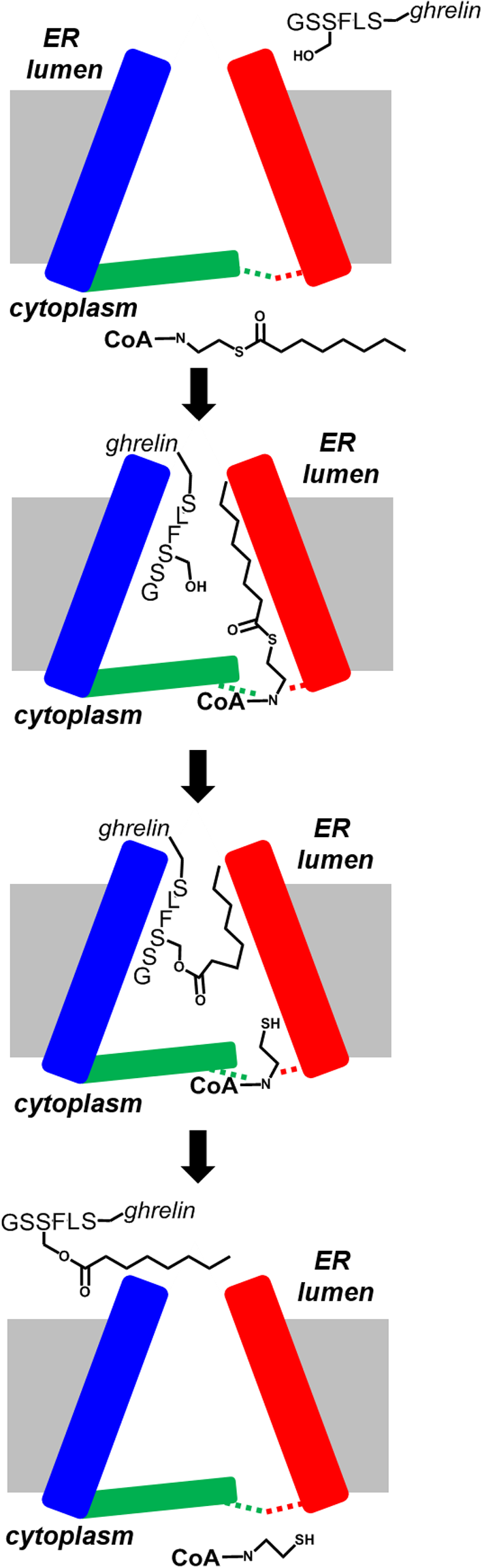
Proposed pathway for transmembrane ghrelin octanoylation by GOAT. Ghrelin (GSSFL-ghrelin) and octanoyl-CoA enter the GOAT internal channel from the ER lumenal pore and cytoplasmic acyl donor binding sites, respectively, followed by acyl transfer to the ghrelin serine side chain hydroxyl. Octanoylated ghrelin dissociates to the ER lumen resulting in the octanoyl chain transiting through the GOAT interior, and coenzyme A is released back to the cytoplasm. The red and blue rectangles represent perimeter helices, the green rectangle represents intramembrane domains forming the cytoplasmic surface of hGOAT, and dotted lines represent binding interactions between the octanoyl-CoA acyl donor and its binding site within hGOAT.

## Acknowledgements

The authors thank Prof. Jason Fridley (Syracuse University) for assistance with statistical analysis and Prof. John Chisholm (Syracuse University) for help with figure generation. The authors also thank Prof. Jinbo Xu (Toyota Technological Institute at Chicago), Prof. Sergey Ovchinnikov (Harvard University), and Prof. Badri Adhikari (University of Missouri–St. Louis) for their valuable advice and suggestions towards computational modeling.

## Author Contributions

MC, FJI, SN, and JLH conceived the project. MC, FJI, TRD, KRMG, RL, SN, and JLH designed the experiments. MC, FJI, TRD, KRMG, RL, MA, NE, MN, and MAS performed the experiments. MC, FJI, TRD, MAS, SN, and JLH analyzed the data. MC, FJI, SN, and JLH wrote the manuscript.

## Abbreviations

GOAT: ghrelin *O*-acyltransferase
MBOAT: membrane bound *O*-acyltransferase
Hhat: Hedgehog acyltransferase
Porcn: Porcupine
hGOAT: human ghrelin *O*-acyltransferase
DltB: D-alanyl transferase DltB
ER: endoplasmic reticulum
TM: transmembrane
IM: intramembrane
RMSD: root-mean-square deviation
CoA: coenzyme-A
WT: wild type
C8: eight carbon acyl group
C12: twelve carbon acyl group
C14: fourteen carbon acyl group

