## Supplemental Data for "Molecular architecture of a membrane-spanning hormone acyltransferase required for metabolic regulation"

### Supplemental Materials

### Materials and Methods

| | |
|---|---|
| **Supplemental File S1.** | Multiple sequence alignment for coevolutionary contact prediction |
| **Supplemental File S2.** | Coevolutionary contact constraints used for hGOAT modeling |
| **Supplemental File S3.** | Alignment of GOAT orthologs |
| **Supplemental Table S1.** | Statistics and parameters for sequence coevolution analysis of hGOAT |
| **Supplemental Table S2.** | hGOAT alanine variant octanoylation activity |
| **Supplemental Table S3.** | Primers for hGOAT mutagenesis |
| **Supplemental Figure S1.** | Flowchart for computational modeling of hGOAT |
| **Supplemental Figure S2.** | Co-evolutionary contact restraints used in hGOAT structural modeling |
| **Supplemental Figure S3.** | Overlay of top 10 lowest energy structures of hGOAT from folding simulations |
| **Supplemental Figure S4**. | Overlay of DltB and hGOAT structures. |
| **Supplemental Figure S5**. | hGOAT variant expression confirmation by anti-Flag Western blotting |
| **Supplemental Figure S6.** | hGOAT alanine variant octanoylation activity mapped onto the hGOAT topology model. |
| **Supplemental Figure S7.** | Acyl donor reactivity of hGOAT alanine variants |
| **Supplemental Figure S8.** | Acyl competition reactions with hGOAT alanine variants |

### Materials and Methods

### Computational methods

*Co-evolutionary contact analysis of hGOAT*

A multiple sequence alignment (MSA) was performed with the hGOAT sequence against the UNIREF90 database utilizing the jackhmmer tool.(*1*) The MSA parameters were set to eight iterative searches (n=8) with an e-value threshold of $1x10^{-40}$. The resulting alignment was filtered to exclude highly similar sequences using the HHfilter tool with 90% identity and 75% sequence coverage cut-offs. This MSA was used as the input for the hmmbuild tool to construct a hidden Markov model (hmm) curated specifically for the MSA,(*2*) which would represent the consensus sequence of hGOAT and its closest homologs. This hmm was then utilized to search against a master database that included uniref100 and metagenome database (metaclust_2018_01) using the hmmsearch tool with a bit score cut-off of 27.(*3, 4*) The resulting MSA was filtered again using the HHfilter tool with 90% identity and 75% sequence coverage against hGOAT. Furthermore, sequences with unidentified amino acids (X, this is to accommodate for RaptorX) and sequence positions with >50% gaps were also filtered from the MSA using trimAL.(*5*) The resulting MSA for hGOAT was primarily used to perform co-evolutionary contact analysis using the RaptorX server and GREMLIN.(*4, 6-8*) The resulting contact maps are provided as Figure 1b (RaptorX) in the text and Supp. Fig S1 (GREMLIN). The resulting MSA had a $M_{eff}$-0.8/$\sqrt{N}$ of 551.7 which is greater than the recommended value of 64 for reliable model prediction using co-evolutionary contacts.(*4, 6*) These contacts were used to guide the hGOAT folding. The MSA analysis and curations were performed using in-house python scripts and conkit python library.(*9*)

*Folding simulations*

The folding simulations were performed in two stages. In both stages, contact restraints were used and the models were iteratively clustered, refined and scored based on their overall backbone energy.

*Stage 1*: In stage 1, a slightly modified version of CONFOLD2 folding routine was used to generate models using either RaptorX contact or GREMLIN contacts.(*10*) The secondary structure predicted from RaptorX-Property was utilized with cross verification by the membrane topology model for mouse GOAT reported by Taylor and coworkers.(*11, 12*) For each of the contact maps, the CONFOLD2 routine was allowed to build 250 structures. The routine is setup in such a way that it will serially utilize 0.1 to 4.0L (where L=435, the length of the amino acid sequence of hGOAT) top contacts at 0.1 intervals from the predicted contact map,(*10*) thus for each contact map 10,000 models were generated. Similarly, the same routine was repeated for GREMLIN contacts. A third run was performed with a hybrid contact map by merging the top 4.0L contacts of RaptorX and GREMLIN after removing all redundant contacts. Thus a sum total of 30,000 models for hGOAT were built and the top 50 models were selected from the pool based on scoring function as mentioned in the CONFOLD2 routine.(*10*) These 50 models were then clustered based on structural similarity and RMSD. The centroid models of the five clusters are once again scored and ranked as 1 to 5 per the CONFOLD2 method.

*Stage 2*: For the second stage of folding simulation, a fragment guided approach was utilized using the ITASSER suite installed on the local computer cluster.(*13, 14*) The long range and medium range contacts observed in the top 5 models from stage 1 are used as the contact restraints in the ITASSER suite (long range contacts are defined as residues >23 amino acids apart in the primary amino acid sequence having a Cβ-Cβ distance of < 8.0 Å; medium range contacts are defined as residues between 12-23 amino acids apart in the primary amino acid sequence having a Cβ-Cβ distance of < 8.0 Å). This meta-contact map contains both the co-evolutionary constraints that guided the stage 1 folding simulations and the energetically favorable contacts that resulted from the folding simulations. Only contacts that are observed in all five models were included in the final folding simulations. The ITASSER simulations were run using L/3, L/2, L and 3L/2 contacts as restraints guiding the folding. The top model from each of these simulations was then clustered and scored based on the scoring function implemented in CONFOLD2.(*10*)

*Refinement and relaxation using molecular dynamics*

The optimized hGOAT model from stage 2 was oriented with respect to a membrane bilayer using the PPM server.(*15*) The calculated hydrophobic thickness of the hGOAT structural model is 25.2 ± 2.4 Å and the tilt angle of 3° relative to the membrane normal vector. The oriented protein was then embedded in a ER-mimetic lipid bilayer (1:1 dipalmitoylphosphatidylcholine(DPPC) : dioleoyl phosphatidylcholine(DOPC)) using the CHARMM- GUI webserver and subject to an all-atom equilibration at 310.15 K in explicit solvent and 150 mM NaCl counter ions.(*16, 17*) The simulation was carried out for 500 ns using GROMACS 2016.4 and the structural deviations were monitored.(*18*) The equilibrated structure was isolated and utilized for prediction of internal channels and docking studies.

*Molecular docking and relaxation of octanoyl-CoA:hGOAT complex*

To build a model of the octanoyl-CoA•hGOAT bound complex, we performed docking using Autodock Vina implemented in the YASARA software suite.(*19, 20*) One hundred docking runs were carried out with the search space defined as 8 Å cube around the residues lining the void space in the model. The top docking pose was further refined with a local search run. The final docked ligand-receptor complex was subject to a short energy minimization and molecular dynamics refinement for 1 ns using the YASARA suite, with the backbone positions being fixed.

### Experimental Methods

*General methods*

Data plotting and curve fitting were carried out with Kaleidagraph (Synergy Software, Reading, PA, USA). Membrane topology schematics were generated using Protter (http://wlab.ethz.ch/protter/start/),(*21*) and structural figures were generated using Chemdraw Prime 15.1 and PyMol. Hexanoyl coenzyme A (hexanoyl-CoA, free acid) (Crystal Chem Inc.), octanoyl coenzyme A (octanoyl-CoA, free acid) (AdventBio), decanoyl coenzyme A (decanoyl-CoA, free acid) (Crystal Chem Inc.), lauroyl coenzyme A (lauroyl-CoA, dodecanoyl-CoA, free acid) (Crystal Chem Inc.), and myristoyl coenzyme

A (myristoyl-CoA, tetradecanoyl-CoA, free acid) (Crystal Chem Inc.) were solubilized to 5 mM in 10 mM Tris-HCl (pH 7.0), aliquoted into low-adhesion microcentrifuge tubes, and stored at −80 °C. Methoxy arachidonyl fluorophosphonate (MAFP) was purchased from Cayman Chemical (Ann Arbor, MI) and solubilized with dimethyl sulfoxide (DMSO). Unlabeled $GSSFLC_{NH2}$ ghrelin peptide was synthesized by Sigma-Genosys (The Woodlands, TX), solubilized in 1:1 acetonitrile:$H_2O$, and stored at -80 °C. Acrylodan (Anaspec) for peptide substrate labeling was solubilized in acetonitrile with the stock concentration determined by absorbance at 393 nm after dilution in methanol ($\varepsilon_{393}$ = 18,483 $M^{-1}cm^{-1}$, per the manufacturer's data sheet). $GSSFLC_{NH2}$ peptide concentrations were determined by reaction of the cysteine thiol with 5,5'-dithiobis(2-nitrobenzoic acid) and absorbance at 412 nm, using $\varepsilon_{412}$ = 14,150 $M^{-1}$ $cm^{-1}$.(*22*)

*Peptide substrate fluorescent labeling*

The $GSSFLC_{NH2}$ peptide substrate used in the hGOAT acylation assay is derived from the N-terminal sequence of ghrelin (GSSFLS) with the C-terminal serine of this peptide (Ser 6) mutated to cysteine to allow chemoselective attachment of an acrylodan fluorophore using our previously reported protocols.(*23, 24*) Acrylodan-labeled peptides were purified by semipreparative reverse phase HPLC (Zorbax Eclipse XDB column, 9.4 x 250 mm) using a gradient mobile phase of 30%-100% acetonitrile in aqueous 0.05 % trifluoroacetic acid (TFA) over 30.2 minutes at a flow rate of 4.2 mL/min. Labeled peptide elution was detected by absorbance at 360 nm, and collected fractions containing the labeled peptides were dried under vacuum at room temperature and resuspended in 1:1 $H_2O$: acetonitrile. $GSSFLC_{NH2}$ labeling was confirmed by MALDI-TOF mass spectrometry

(Bruker Autoflex III) using a matrix containing sinapinic acid in 0.1 % TFA and 50 mM ammonium phosphate. The concentration of acrylodan labeled $GSSFLC_{NH2}$ was calculated using absorbance of acrylodan at 360 nm (ε= 13,300 $M^{-1}cm^{-1}$) per previous reports.(*24, 25*)

*Construction of hGOAT mutants*

PCR primers for site-directed mutagenesis were designed from our previously reported hGOAT expression construct (Table S3).(*24*) This construct was commercially synthesized by Integrated DNA Technologies (Coralville, IA) containing a C-terminal FLAG epitope tag, a polyhistidine ($His_6$) tag, and 3x human influenza hemagglutinin (HA) tags appended downstream of a TEV protease site.(*26*) Primers were dissolved in $H_2O$ and concentrations were measured by UV absorbance at 260 nm. PCR mutagenesis reactions (50 µL total volume) contained the following components: 10x Pfu reaction buffer (5 µL), template plasmid DNA (10 ng), forward primer (125 ng), reverse primer (125 ng), dNTPs (1 µL, 1 mM stock), and Pfu Turbo DNA polymerase (1 µL, Agilent). The thermocycler program for PCR mutagenesis proceeded as follows: initial denaturation (95 °C, 1 min); 18 cycles of denaturation (95 °C, 50 sec); annealing (60 °C, 50 sec); extension (68 °C, 12 min); and final extension (68 °C, 12 min). Reaction were digested with *Dpn*I (New England BioLabs, R0176S, 10 units) at 37 °C for 2 hours. After *Dpn*I digestion, the PCR reaction mixture (5 µL) was transformed into a 100 µL aliquot of chemically competent Z-competent DH5α cells (Zymo Research) followed by incubation on ice for 30 minutes. Transformed bacteria were spread on LB-ampicillin plates (100 µg/mL) and incubated at 37 °C overnight. Following overnight incubation at 37 °C, single colonies were inoculated into LB media (5 mL) containing ampicillin (100 μg/mL) in sterile culture tubes. These cultures were incubated overnight at 37 °C with shaking (225 rpm). Following overnight growth, plasmids were purified from the saturated cultures using EZ-10 Spin Column Plasmid DNA kit (Bio Basic Inc.) per manufacturer's instructions. Single site mutations were verified by DNA sequencing (Genewiz).

*Expression and enrichment of hGOAT in membrane protein fractions*

hGOAT wildtype and mutants were expressed in insect (Sf9) cell membrane fractions using previously published procedures. (*24, 27, 28*)

*Membrane fraction enrichment*

Sf9 insect cell expression cultures were transferred to 50 mL conical tubes and harvested by centrifugation (500 x *g*, 5 min, 4°C); the resulting supernatant was discarded. The cell pellet was resuspended in 2.5 mL lysis buffer (50 mM Tris-HCl pH 7.0, 150 mM sodium chloride (NaCl), 1 mM sodium ethylenediamine (NaEDTA), 1 mM dithiothreitol (DTT), 0.01 μg/mL pepstatin A, 10 μM bis(4-nitrophenyl)phosphate, and 1 mini Roche protease inhibitor cocktail tablet/10 mL). The cells were then lysed with a 7 mL dounce homogenizer on ice, using 20 strokes with a small pestle and 20 strokes with a large pestle. Intact cells and cell debris were removed by centrifugation (3000 x *g*, 10 min, 4°C). The supernatant was transferred to a pre-tared ultracentrifuge tube and the membrane fraction was isolated by centrifugation (100,000 x g, 1 hour, 4°C). The resulting supernatant was discarded, and the pellet was solubilized in 50 mM HEPES (pH

7.0) to reach 4 μL/mg of microsomal pellet. Membrane fraction suspensions are distributed into 80 μL aliquots in low-adhesion tubes and stored at -80°C. (*24*).

*hGOAT expression analysis by anti-FLAG Western Blot*

hGOAT membrane fractions were thawed on ice and homogenized by passing through an 18-gauge needle ten times. Membrane fraction protein concentrations were determined by Bradford assay using the Quick Start Bradford 1X Dye Reagent (BioRad). Samples for polyacrylamide gel analysis and Western blotting were prepared containing 50 μg membrane fraction protein from Sf9 cells transfected with hGOAT, 1x sample buffer (0.33 M Tris HCl, pH 6.8, 0.1 M SDS, 14% glycerol, and 0.5 M DTT) and 50 mM HEPES pH 7.0 in a total volume of 45 μL. Protein samples were heated to 50.2 °C for 5 minutes and then incubated at room temperature for 15 min prior to gel loading. Samples were loaded onto a 12 % Tris-glycine SDS-polyacrylamide gel and run at 110 V for 1.5 hrs. Each gel contained an empty vector (EV) microsomal protein as negative control and amino-terminal FLAG-BAP Fusion protein as a positive control (Millipore Sigma, P7582-100UG, 1:150 dilution, 30 μL total volume).

Following electrophoretic separation, proteins were transferred to a PVDF membrane for Western blotting (BioRad, Trans-Blot turbo RTA transfer kit), with the PVDF membrane activated by immersion in methanol for 30 seconds followed by equilibration in transfer buffer (20% v/v methanol, 48 mM Tris base, 39 mM glycine and 0.034% v/v SDS) prior to transfer. Proteins were transferred to the membrane for 30 minutes at 1.3 A / 25 V using the transfer kit per manufacturer's instructions. Following electroblotting, the PVDF membrane was blocked for 4 hours in 10% v/v nonfat milk in TBST buffer (Tris buffered saline (TBS, 0.05M Tris and 0.14M NaCl) with 0.1% v/v Tween 20). The membrane was then probed with a Flag antibody (HRP-conjugated DYKDDDDK Tag Antibody, Invitrogen catalog number PA1-984B-HRP, 1:1000 dilution, 10 mL total volume) in 5% nonfat milk in TBST buffer overnight at 4 °C. The membrane was washed with TBST (6 x 5 mL) and treated with West Pico Chemiluminescent substrate-imaging reagent (Thermo Scientific), followed by imaging on a ChemiDoc XRS+ gel documentation system (BioRad).

*hGOAT activity assay - standard reaction conditions*

Membrane fractions of hGOAT expressed from Sf9 cells were thawed on ice and homogenized by passing through an 18-gauge needle ten times. hGOAT activity assays under standard conditions were performed with 50 μg of membrane protein, 1.5 μM fluorescent peptide substrate, 300 μM octanoyl-CoA, 1 μM MAFP, and 50 μM HEPES pH 7.0 in a total volume of 50 μL. All components except for the peptide and acyl-CoA substrates were incubated at room temperature for 30 minutes prior to reaction initiation by addition of peptide and acyl-CoA substrates. Reactions were incubated at room temperature for 2 hour in the dark and then stopped by addition of 50 μL of 20% acetic acid in isopropanol. Reaction solutions were clarified by protein precipitation with 16.7 μL of 20% trichloroacetic acid followed by centrifugation (1,000 x *g*, 2 min). The resulting supernatant was then analyzed by reverse phase HPLC. hGOAT assay samples were analyzed on an Agilent 1260 HPLC with a C18 reverse phase HPLC column (Zorbax Eclipse XDB, 4.6 x 150 mm) using a gradient of 30% acetonitrile in 0.05 % aqueous TFA to 100 % acetonitrile over 30.2 minutes. Fluorescent peptide substrate and acylated products were detected by UV absorbance at 360 nm and fluorescence ($\lambda_{ex}$ 360 nm, $\lambda_{em}$ 485 nm), with the peptide substrate eluting with a retention time of 5-6 minutes and the octanoylated peptide eluting with a retention time of 11-12 minutes. Chromatogram analysis and peak integration was performed using Chemstation for LC (Agilent Technologies).(*24*) Product conversion was calculated by dividing the integrated fluorescence for the product peak by the total integrated peptide fluorescence (substrate and product) in each run. Percent activity for each hGOAT mutant was calculated by normalizing the product conversion for the mutant to that of wild type hGOAT in a reaction run in parallel on the same day using the same reagents.

*Statistical analysis of hGOAT alanine variant reactivity.*

Each alanine mutation site was assigned as "void contacting" or "non-void contacting" by inspection of the hGOAT structural model. The two populations were compared using a Wilcoxon signed-rank test (n=42, test statistic W=294.5) and yielded a p-value = 0.02978 against the null hypothesis of both populations of alanine mutations being equally likely to yield reduced hGOAT activity. The statistical test was executed using the script below executed in R:(*29*)

```
str(dat)
median(dat$Y)
   #overall median is 13.25
sum(dat$group)
   #21 values in interest group
median(dat$Y[dat$group==1])
   #interest group median is 4.41
median(dat$Y[dat$group==0])
   #median of other group is 33.72
actual.diff = median(dat$Y[dat$group==0]) - median(dat$Y[dat$group==1])
```

```
    #difference of group medians is 29.3

#are the group medians significantly different?
#procedure: randomly shuffle group membership vector, calculate median difference as
test statistic

Nperm = 10000 #number of random permutations
test.stat = rep(0,Nperm) #holding vector
for(i in 1:Nperm) {
    rgroup = sample(dat$group)
    test.stat[i] = median(dat$Y[rgroup==0]) - median(dat$Y[rgroup==1])
}
hist(test.stat); abline(v=actual.diff,lwd=3,col="red")
length(test.stat[test.stat>actual.diff])/Nperm #P value of null hypothesis of no difference
    #P ~ 0.04 (one-tailed)

#compare to t.test of mean difference assuming unequal variances, one-tailed
t.test(dat$Y[dat$group==0],dat$Y[dat$group==1],alternative="greater")
    #P=0.013
#compare to wilcoxon rank sum test (nonparameteric)
wilcox.test(dat$Y[dat$group==0],dat$Y[dat$group==1],alternative="greater")
    #P=0.030
```

*Single acyl donor reactivity assay*

To determine the reactivity of hGOAT variants with different length acyl donors, hGOAT activity was measured as described above in the presence of 100 μM of a single acyl donor (octanoyl-CoA, lauryl (dodecanoyl)-CoA, or myristoyl (tetradecanoyl)-CoA). Reactions were initiated and analyzed as described above, with product conversions calculated as described above. Characteristic retention times for each acylated form of the peptide substrate provided confirmation of the nature of the attached acyl chain, with dodecanoyl-GSSFLC$_{AcDan}$ eluting at ~17 minutes and tetradecanoyl- GSSFLC$_{AcDan}$ eluting at ~19 minutes under the standard HPLC gradient for hGOAT assay analysis. For each acyl donor, relative activity was calculated normalized to the highest activity observed across the panel of wildtype hGOAT and hGOAT variants.

*Acyl donor competition assay*

To determine the relative preference of each hGOAT variant for acyl donors ranging from six to twelve carbons, hGOAT activity was measured as described above in the presence of 100 μM each of four potential acyl donors (hexanoyl-CoA, octanoyl-CoA, decanoyl-CoA, and lauryl (dodecanoyl)-CoA). Reactions were performed as described above, with analysis by reverse-phase HPLC with fluorescence detection of the acrylodan fluorophore. Each potential product peak was assigned by retention time compared to a standard reaction containing only one acyl donor for each potential product. Competition experiments including myristoyl (tetradecanoyl)-CoA) were unsuccessful, potentially due to low critical micelle concentration (CMC) for this acyl donor lying near 100 μM.(*30*)

**Supplemental Table S1.** Statistics and parameters for sequence coevolution analysis of hGOAT.

<table>
<tr><th colspan="3">Human ghrelin O-acyltransferase</th></tr>
<tr><td>Query sequence</td><td colspan="2">Homo sapiens MBOAT (UniProt Q96T53)</td></tr>
<tr><td>Alignment window</td><td colspan="2">1-435 (15-420)</td></tr>
<tr><td>Alignment</td><td colspan="2">Jackhmmer, HHblits, hhmsearch<br>metaclust_2018_01, Uniref100.</td></tr>
<tr><td>Filtering</td><td colspan="2">90% identity 75 % coverage and filtered columns with >50% gaps</td></tr>
<tr><td rowspan="12">$M_{eff}$</td><td colspan="2">$M_{eff}$ at various identity cut-off of the MSA</td></tr>
<tr><td>Identity</td><td>Meff</td></tr>
<tr><td>50.0%</td><td>726</td></tr>
<tr><td>55.0%</td><td>1450</td></tr>
<tr><td>60.0%</td><td>2714</td></tr>
<tr><td>65.0%</td><td>4516</td></tr>
<tr><td>70.0%</td><td>6809</td></tr>
<tr><td>75.0%</td><td>9193</td></tr>
<tr><td>80.0%</td><td>11507</td></tr>
<tr><td>85.0%</td><td>13701</td></tr>
<tr><td>90.0%</td><td>16114</td></tr>
<tr><td>95.0%</td><td>16555</td></tr>
<tr><td>$M_{eff}$-0.8/$\sqrt{N}$</td><td colspan="2">551.7</td></tr>
<tr><td rowspan="2">Sequences / L</td><td colspan="2">>45; Sequence diversity ($\sqrt{N}/L$): 0.296</td></tr>
<tr><td colspan="2">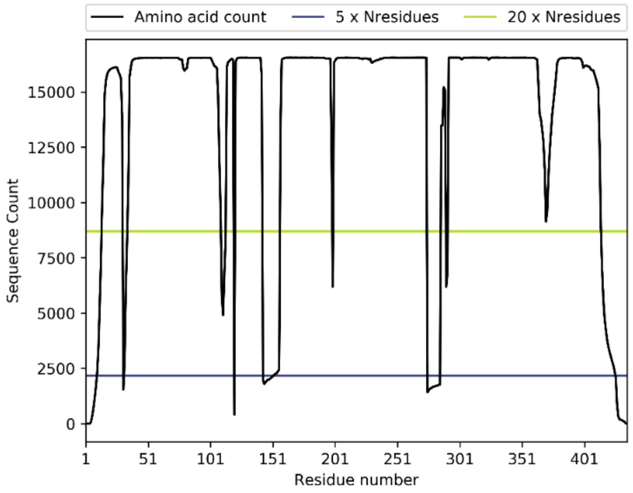
</td></tr>
<tr><td>Coevolution algorithm</td><td colspan="2">Raptor-X (machine Learning) and Gremlin (Markov Random Field pseudo-likelihood maximization)</td></tr>
<tr><td>Significance cut-off</td><td colspan="2">Of the 3L/2 direct Co-evolutionary contacts predicted by GREMLIN 93% of the contacts were observed at or below 10Å in the final structure; 86% of the contacts were observed at or below 8Å</td></tr>
<tr><td>Comparison with DltB</td><td colspan="2">Distant homolog within the same family<br>CA-RMSD of aligned 98 residues is 2.23 Å</td></tr>
</table>

**Supplemental Table S2.** hGOAT alanine variant octanoylation activity with the $GSSFLC_{AcDan}$ peptide substrate under standard reaction conditions. Reactions were performed, analyzed, and normalized as described in the Experimental Methods. Errors reflect the standard deviation of a minimum of three independent experimental trials.

| **Active Variants (100 - 33% of WT Activity)** | | | **Impaired Variants (33 - 1% of WT Activity)** | | | **Inactive Variants (Undetectable Activity)** | |
|---|---|---|---|---|---|---|---|
| Variant | % Activity | | Variant | % Activity | | Variant | % Activity |
| S179A | 60 ± 8 | | L96A | 27 ± 4 | | H98A | 0 |
| Q191A | 92 ± 30 | | L135A | 8.3 ± 0.9 | | S132A | 0 |
| H258A | 56 ± 8 | | C181A | 20 ± 1 | | Y168A | 0 |
| D262A | 61 ± 12 | | S182A | 2.4 ± 0.7 | | L180A | 0 |
| E281A | 74 ± 3 | | F183A | 2.4 ± 0.1 | | D234A | 0 |
| D289A | 58 ± 3 | | Y255A | 21 ± 1 | | C235A | 0 |
| E294A | 43 ± 3 | | D287A | 6 ± 2 | | D263A | 0 |
| H297A | 56 ± 4 | | S300A | 4 ± 1 | | E282A | 0 |
| Q320A | 94 ± 2 | | F302A | 27.4 ± 0.9 | | Y284A | 0 |
| H341A | 44 ± 7 | | F331A | 18.2 ± 0.6 | | R304A | 0 |
| F348A | 67 ± 5 | | W351A | 4.7 ± 0.9 | | W306A | 0 |
| F364A | 94 ± 13 | | H362A | 29 ± 7 | | N307A | 0 |
| R370A | 62 ± 10 | | | | | R315A | 0 |
| L381A | 128 ± 5 | | | | | H338A | 0 |
| L423A | 106 ± 19 | | | | | D358A | 0 |

**Supplemental Table S3.** Primers for hGOAT alanine mutagenesis

| Mutation | Primers |
|---|---|
| L96A | Forward: CAAACAGCTTGTCACCTGGGACTCCAC |
| | Reverse: GTGGAGTCCCAGGTGACAAGCTGTTTG |
| H98A | Forward: GCTGGCAAACACTGTGTGCTCTGGGACTCCACTAC |
| | Reverse: GTAGTGGAGTCCCAGAGCACACAGTGTTTGCCAGC |
| S132A | Forward: CTCAGCGCGTGACAGCTTTGTCTCTGGACATC |
| | Reverse: GATGTCCAGAGACAAAGCTGTCACGCGCTGAG |
| L135A | Forward: GTGACATCCTTGTCTGCTGACATCTGTGAGGGC |
| | Reverse: GCCCTCACAGATGTCAGCAGACAAGGATGTCAC |
| Y168A | Forward: CTGCCCTACTTCTCCGCTCTGCTCTTCTTCCC |
| | Reverse: GGGAAGAAGAGCAGAGCGGAGAAGTAGGGCAG |
| S179A | Forward: GCCTTGCTGGGAGGTGCTCTCTGTTCATTCCAG |
| | Reverse: CTGGAATGAACAGAGAGCACCTCCCAGCAAGGC |
| L180A | Forward: GCTGGGAGGTAGCGCTTGTTCATTCCAGCG |
| | Reverse: CGCTGGAATGAACAAGCGCTACCTCCCAGC |
| C181A | Forward: CTGGGAGGTAGCCTCGCTTCATTCCAGC |
| | Reverse: GCTGGAATGAAGCGAGGCTACCTCCCAG |
| S182A | Forward: GAGGTAGCCTCTGTGCTTTCCAGCGTTTCCAAG |
| | Reverse: CTTGGAAACGCTGGAAAGCACAGAGGCTACCTC |
| F183A | Forward: GGTAGCCTCTGTTCAGCTCAGCGTTTCCAAGC |
| | Reverse: GCTTGGAAACGCTGAGCTGAACAGAGGCTACC |
| Q191A | Forward: CAAGCTAGAGTCGCTGGATCGTCCGCTC |
| | Reverse: GAGCGGACGATCCAGCGACTCTAGCTTG |
| D234A | Forward: GCCGGACTGACTGCTTGCCAGCAATTCGAATG |
| | Reverse: CATTCGAATTGCTGGCAAGCAGTCAGTCCGGC |
| C235A | Forward: GGAGCCGGACTGACTGATGCCCAGCAATTCGAATG |
| | Reverse: CATTCGAATTGCTGGGCATCAGTCAGTCCGGCTCC |
| E239A | Forward: GATTGCCAGCAATTCGCTTGTATCTACGTTG |
| | Reverse: CAACGTAGATACAAGCGAATTGCTGGCAATC |
| Y255A | Forward: GGTTGTTCAAGCTGACCGCTTACTCACACTGGATC |
| | Reverse: GATCCAGTGTGAGTAAGCGGTCAGCTTGAACAACC |
| H258A | Forward: GCTGACCTACTACTCAGCCTGGATCCTCGACG |
| | Reverse: CGTCGAGGATCCAGGCTGAGTAGTAGGTCAGC |
| D262A | Forward: CACACTGGATCCTCGCCGATTCGCTCTTGC |
| | Reverse: GCAAGAGCGAATCGGCGAGGATCCAGTGTG |
| D263A | Forward: CTGGATCCTCGACGCTTCGCTCTTGCACG |
| | Reverse: CGTGCAAGAGCGAAGCGTCGAGGATCCAG |

| | |
|---|---|
| E281A | Forward: GACAGTCACCAGGAGCGGAAGGTTACGTTCC |
| | Reverse: GGAACGTAACCTTCCGCTCCTGGTGACTGTC |
| E282A | Forward: GTCACCAGGAGAGGCAGGTTACGTTCCTG |
| | Reverse: CAGGAACGTAACCTGCCTCTCCTGGTGAC |
| Y284A | Forward: CCAGGAGAGGAAGGTGCTGTTCCTGACGCTGATATC |
| | Reverse: GATATCAGCGTCAGGAACAGCACCTTCCTCTCCTGG |
| D287A | Forward: GGTTACGTTCCTGCCGCTGATATCTGGACC |
| | Reverse: GGTCCAGATATCAGCGGCAGGAACGTAACC |
| D289A | Forward: CGTTCCTGACGCTGCTATCTGGACCCTGG |
| | Reverse: CCAGGGTCCAGATAGCAGCGTCAGGAACG |
| E294A | Forward: GATATCTGGACCCTGGCAAGGACTCACAGAATC |
| | Reverse: GATTCTGTGAGTCCTTGCCAGGGTCCAGATATC |
| H297A | Forward: CCCTGGAAAGGACTGCCAGAATCTCGGTCTTC |
| | Reverse: GAAGACCGAGATTCTGGCAGTCCTTTCCAGGG |
| S300A | Forward: GGACTCACAGAATCGCTGTCTTCTCCCGTAAG |
| | Reverse: CTTACGGGAGAAGACAGCGATTCTGTGAGTCC |
| F302A | Forward: CACAGAATCTCGGTCGCTTCCCGTAAGTGGAAC |
| | Reverse: GTTCCACTTACGGGAAGCGACCGAGATTCTGTG |
| R304A | Forward: GAATCTCGGTCTTCTCCGCTAAGTGGAACCAAAGC |
| | Reverse: GCTTTGGTTCCACTTAGCGGAGAAGACCGAGATTC |
| W306A | Forward: GTCTTCTCCCGTAAGGCTAACCAAAGCACTGCTC |
| | Reverse: GAGCAGTGCTTTGGTTAGCCTTACGGGAGAAGAC |
| N307A | Forward: CTCCCGTAAGTGGGCCCAAAGCACTGCTCGC |
| | Reverse: GCGAGCAGTGCTTTGGGCCCACTTACGGGAG |
| R315A | Forward: CTGCTCGCTGGCTCGCTCGCTTGGTGTTCC |
| | Reverse: GGAACACCAAGCGAGCGAGCCAGCGAGCAG |
| Q320A | Forward: CGCTTGGTGTTCGCTCACAGCCGTGCTTG |
| | Reverse: CAAGCACGGCTGTGAGCGAACACCAAGCG |
| F331A | Forward: CCACTGCTCCAAACAGCTGCTTTCTCAGCTTGG |
| | Reverse: CCAAGCTGAGAAAGCAGCTGTTTGGAGCAGTGG |
| H338A | Forward: CAGCTTGGTGGGCCGGACTGCACCCTGG |
| | Reverse: CCAGGGTGCAGTCCGGCCCACCAAGCTG |
| H341A | Forward: GCACGGACTGGCCCCTGGACAGGTTTTCGG |
| | Reverse: CCGAAAACCTGTCCAGGGGCCAGTCCGTGC |
| F348A | Forward: GACAGGTTTTCGGTGCTGTGTGCTGGGCTGTTATG |
| | Reverse: CATAACAGCCCAGCACACAGCACCGAAAACCTGTC |
| W351A | Forward: CGGTTTCGTGTGCGCTGCTGTTATGGTGGAG |
| | Reverse: CTCCACCATAACAGCAGCGCACACGAAACCG |
| D358A | Forward: GTTATGGTGGAGGCCGCCTACCTGATCCAC |
| | Reverse: GTGGATCAGGTAGGCGGCCTCCACCATAAC |

| | |
|---|---|
| H362A | Forward: GCCGACTACCTGATCGCCTCCTTCGCTAACGAG |
| | Reverse: CTCGTTAGCGAAGGAGGCGATCAGGTAGTCGGC |
| R370A | Forward: GCTAACGAGTTCATCGCTTCTTGGCCAATGAGG |
| | Reverse: CCTCATTGGCCAAGAAGCGATGAACTCGTTAGC |
| L381A | Forward: CTTCTACAGAACAGCTACCTGGGCCCAC |
| | Reverse: GTGGGCCCAGGTAGCTGTTCTGTAGAAG |
| L395A | Forward: GCTTACATCATGGCTGCCGTCGAGGTTAG |
| | Reverse: CTAACCTCGACGGCAGCCATGATGTAAGC |
| L423A | Forward: GATGGTTTACTGTATCGCTTTGCTGCTCTTG |
| | Reverse: CAAGAGCAGCAAAGCGATACAGTAAACCATC |

**Supplemental Figure S1.** Flowchart for computational modeling of hGOAT

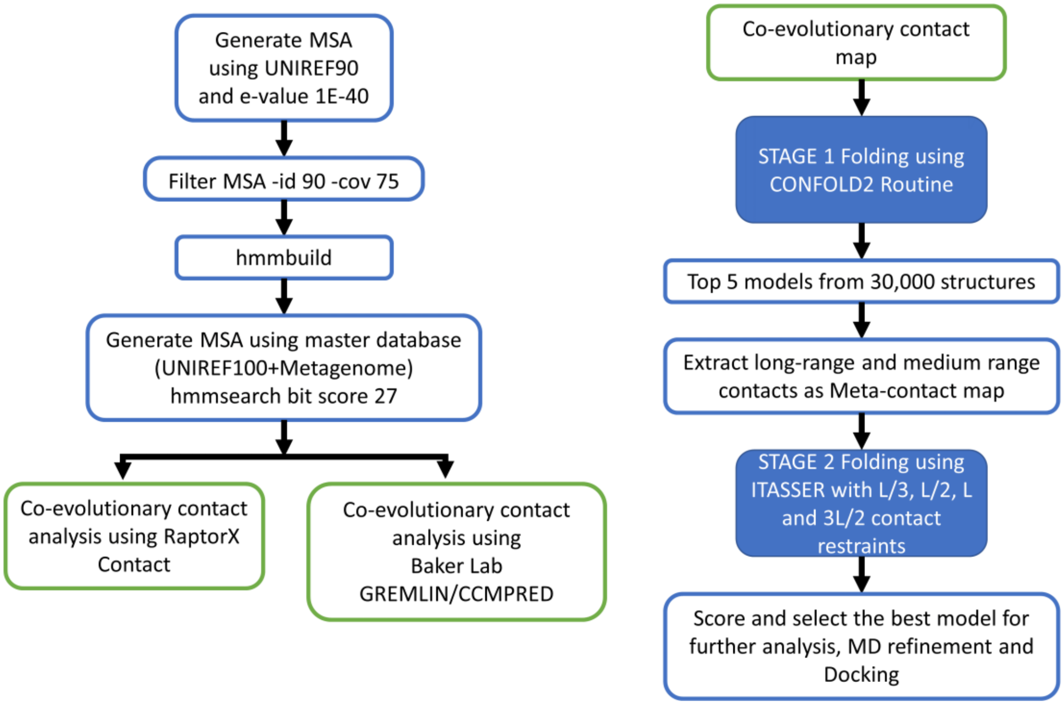

**Supplemental Figure S2.** Co-evolutionary contact restraints used in hGOAT structural modeling. a) Contact map for hGOAT showing the probability for a coevolutionary contact from GREMLIN analysis. b) Heat map of co-evolutionary contact constraints mapped onto hGOAT structure (red: highest number of co-evolutionary constraints per residue (8); white: no co-evolutionary constraints)

a)

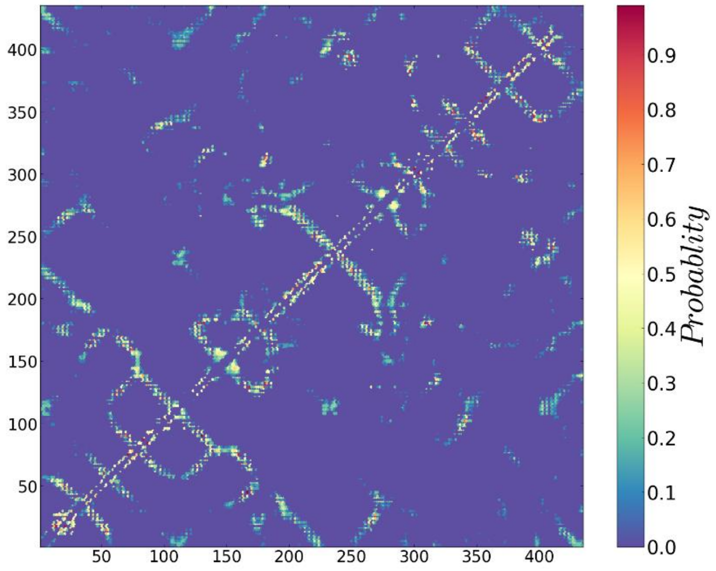

b)

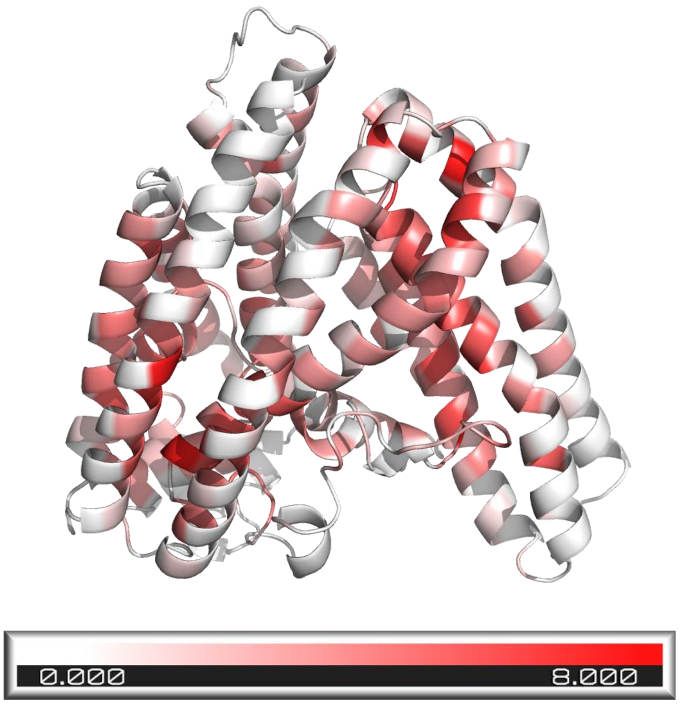

**Supplemental Figure S3.** Overlay of top 10 lowest energy structures of hGOAT from folding simulations.

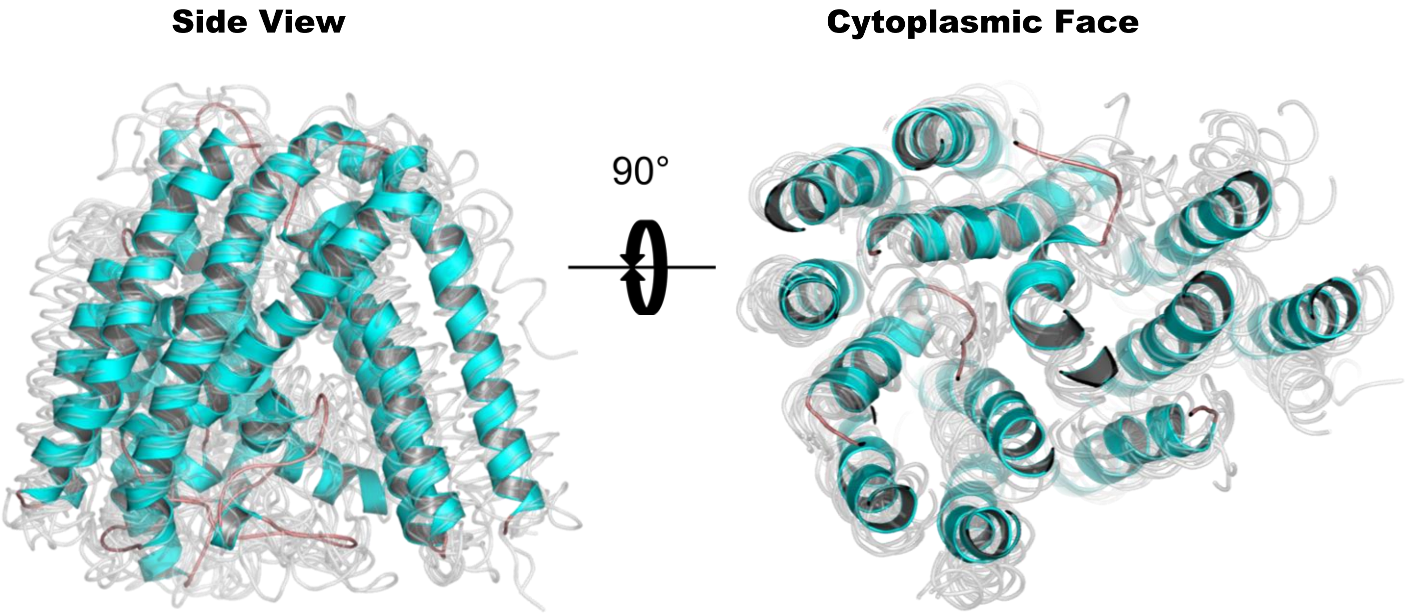

**Supplemental Figure S4**. Overlay of DltB and hGOAT structures. a) Structure of DltB (PDB ID 6BUG:C). b) Overlay of DltB (purple) and hGOAT (teal) structures with side view on left and cytoplasmic face view on right.

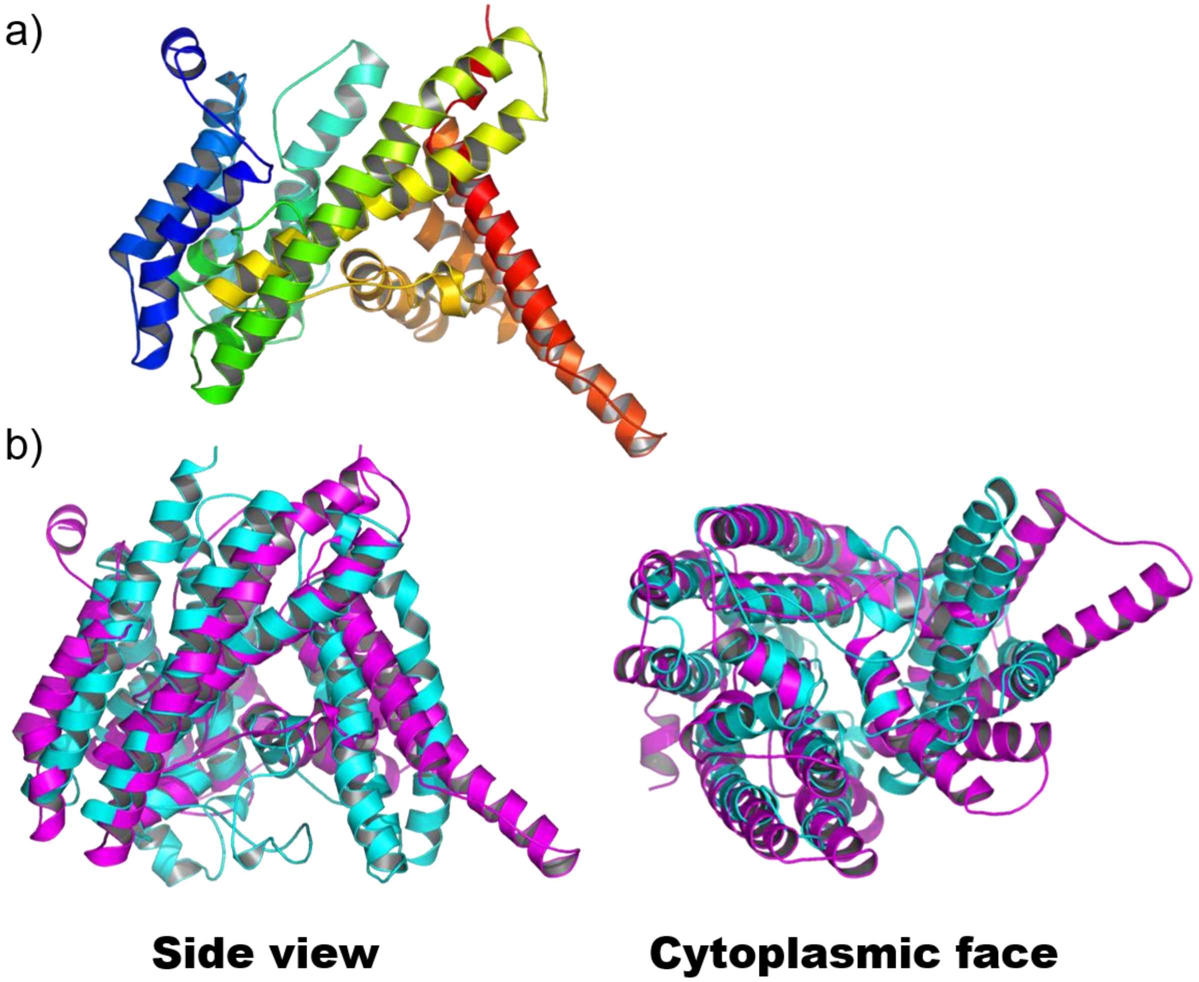

**Supplemental Figure S5**. hGOAT variant expression confirmation by anti-Flag Western blotting. Western blots were performed as described in Materials and Method. WT, wild type hGOAT; EV, empty vector baculoviral expression; Flag, Flag-BAP fusion protein positive control.

| Variant | Blot | | Variant | Blot | | Variant | Blot | | Variant | Blot |
|---|---|---|---|---|---|---|---|---|---|---|
| L96A | b | | D234A | b | | E294A | a | | H341A | f |
| H98A | a | | C235A | b | | H297A | e | | F348A | f |
| S132A | c | | Y255A | c | | S300A | e | | W351A | g |
| L135A | c | | H258A | d | | F302A | e | | D358A | d |
| Y168A | a | | D262A | d | | R304A | e | | H362A | g |
| S179A | a | | D263A | b | | W306A | c | | F364A | g |
| L180A | a | | E281A | d | | N307A | e | | R370A | e |
| C181A | a | | E282A | d | | R315A | c | | L381A | g |
| S182A | b | | Y284A | a | | Q320A | f | | L423A | g |
| F183A | c | | D287A | d | | F331A | f | | | |
| Q191A | b | | D289A | d | | H338A | f | | | |

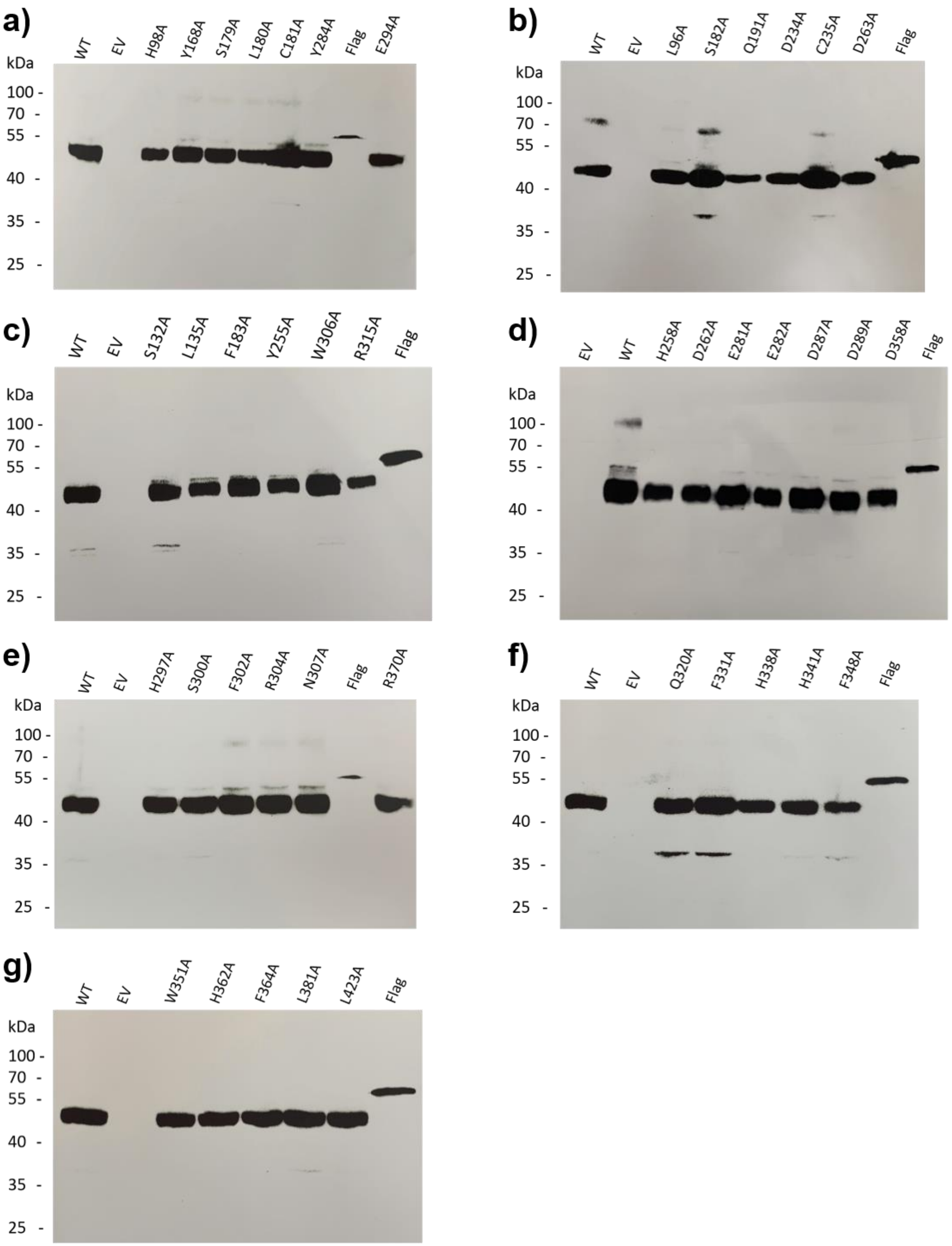
a)
WT EV H98A Y168A S179A L180A C181A Y284A Flag E294A
kDa
100 70 55 40 35 25
b)
WT EV L96A S182A Q191A D234A C235A D263A Flag
kDa
100 70 55 40 35 25
c)
WT EV S132A L135A F183A Y255A W306A R315A Flag
kDa
100 70 55 40 35 25
d)
EV WT H258A D262A E281A E282A D287A D289A D358A Flag
kDa
100 70 55 40 35 25
e)
WT EV H297A S300A F302A R304A N307A Flag R370A
kDa
100 70 55 40 35 25
f)
WT EV Q320A F331A H338A H341A F348A Flag
kDa
100 70 55 40 35 25
g)
WT EV W351A H362A F364A L381A L423A Flag
kDa
100 70 55 40 35 25

**Supplemental Figure S6.** hGOAT alanine variant octanoylation activity mapped onto the hGOAT topology model. Blue squares, alanine variants with octanoylation activity within 3-fold of WT hGOAT; purple diamonds, alanine variants with impaired octanoylation activity (>3-fold loss compared to WT hGOAT); red squares, inactive alanine variants.

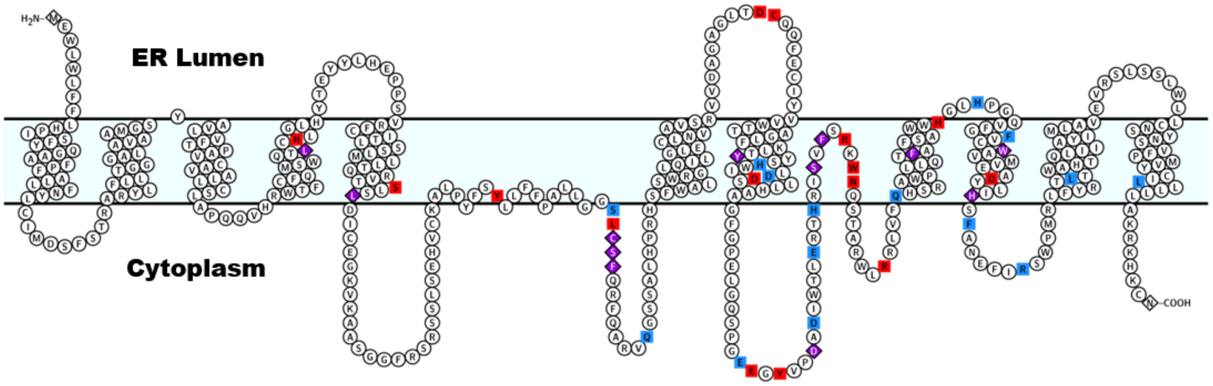

**Supplemental Figure S7.** Acyl donor reactivity with WT hGOAT and hGOAT alanine variants with C8-CoA (black), C12-CoA (purple), and C14-CoA (green) acyl donors. Single acyl donor reactions were performed, analyzed, and normalized as described in the Experimental Methods. Error bars reflect the standard deviation of a minimum of three independent experimental trials.

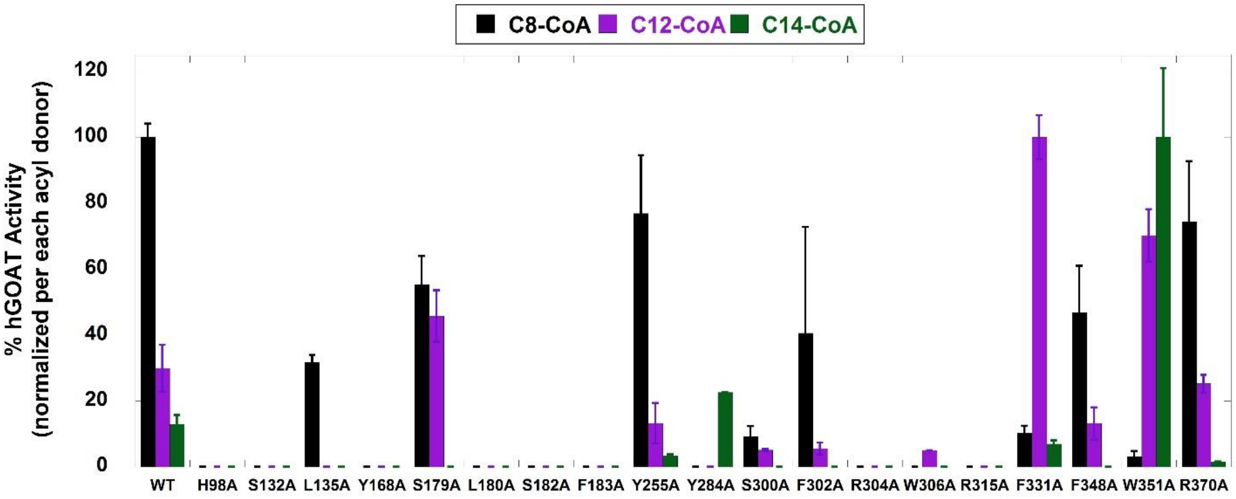

**Supplemental Figure S8.** Acyl competition reactions with WT hGOAT and hGOAT alanine variants with C6-CoA, C8-CoA, C10-CoA, and C12-CoA. Reactions were performed, analyzed, and normalized as described in the Experimental Methods. a) WT; b) H98A; c) L135A; d) Y168A; e) S179A; f) L180A; g) S182A; h)F183A; i) Y255A, j) Y284A; k) S300A; l) F302A; m) W306A; n) R315A; o) F331A; p) F348A; q) W351A; r) R370A.

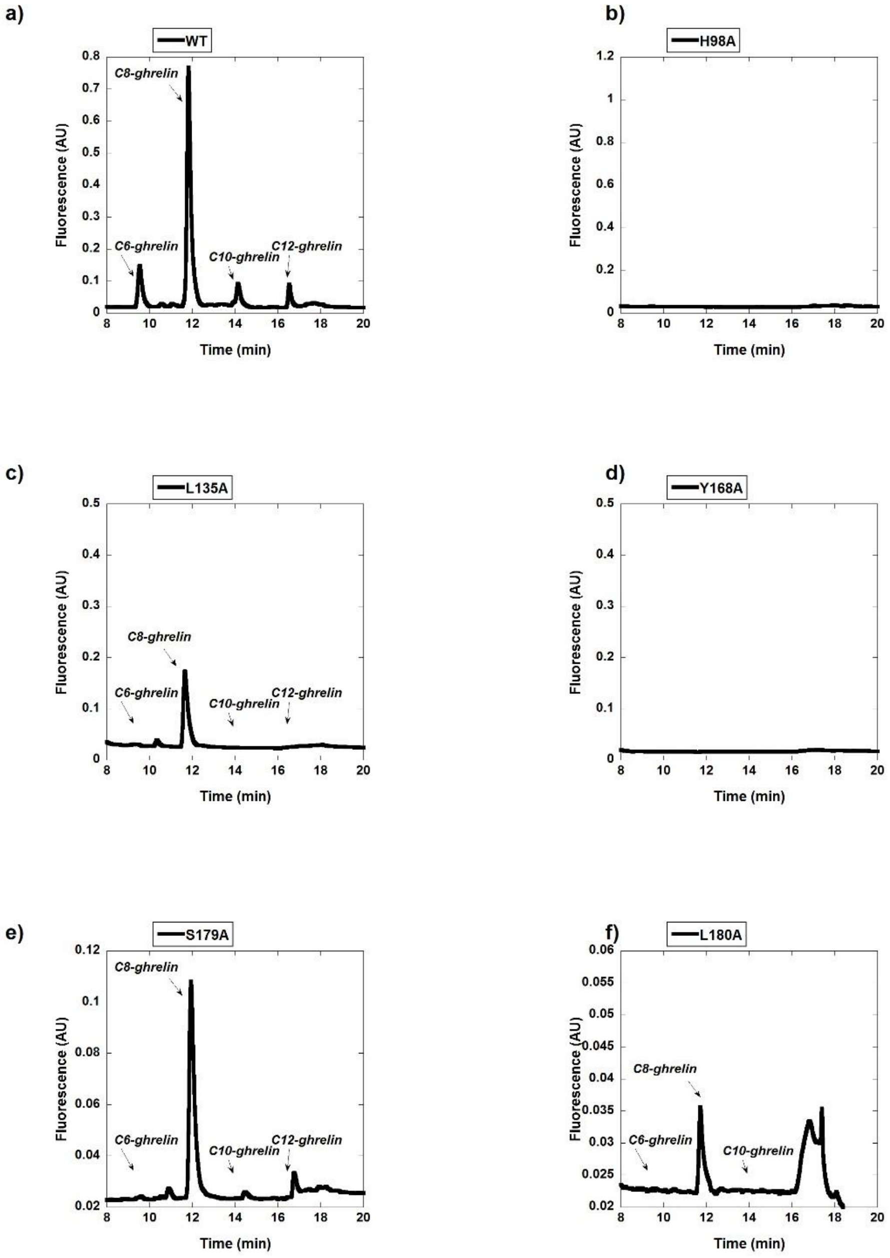

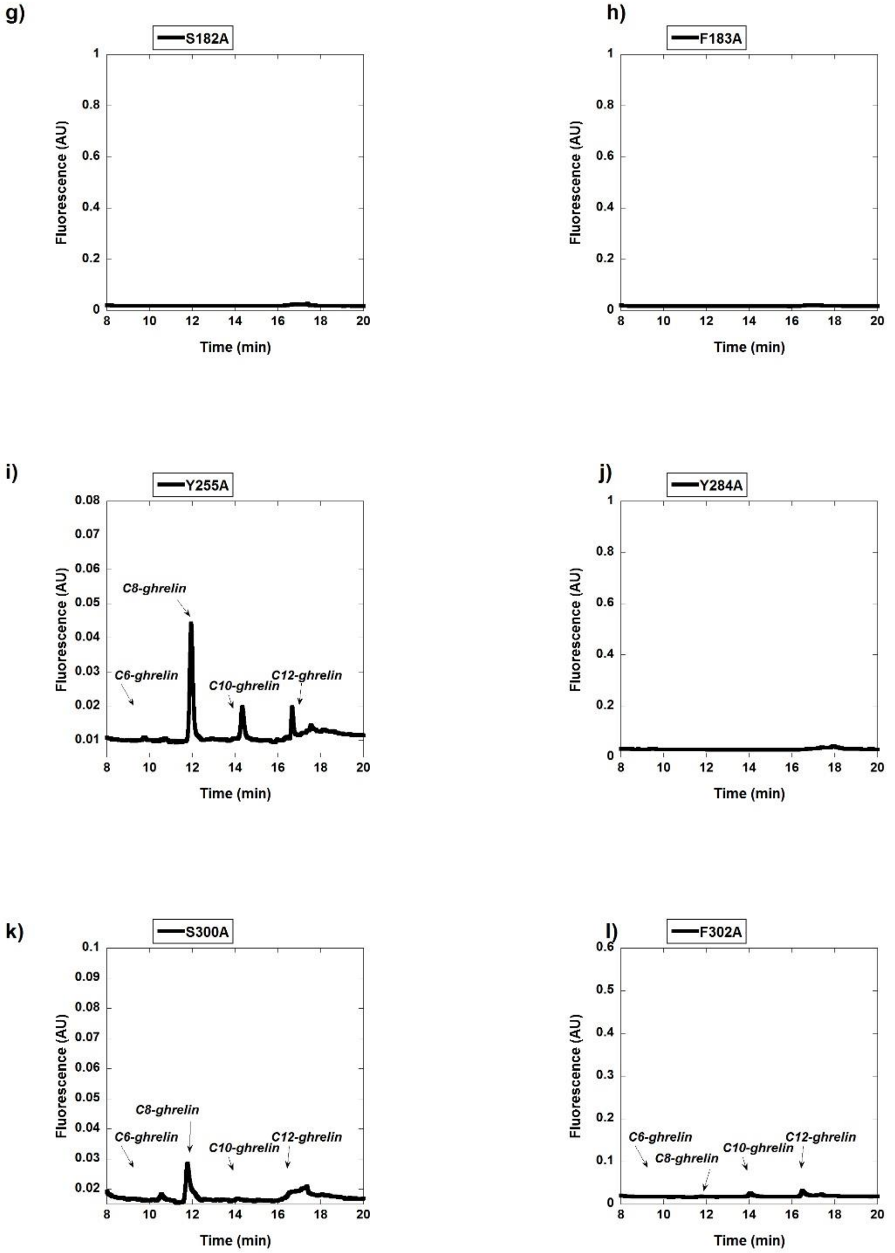
g)
S182A
Fluorescence (AU)
Time (min)
h)
F183A
Fluorescence (AU)
Time (min)
i)
Y255A
C8-ghrelin
C6-ghrelin
C10-ghrelin
C12-ghrelin
Fluorescence (AU)
Time (min)
j)
Y284A
Fluorescence (AU)
Time (min)
k)
S300A
C8-ghrelin
C6-ghrelin
C10-ghrelin
C12-ghrelin
Fluorescence (AU)
Time (min)
l)
F302A
C6-ghrelin
C8-ghrelin
C10-ghrelin
C12-ghrelin
Fluorescence (AU)
Time (min)

m)

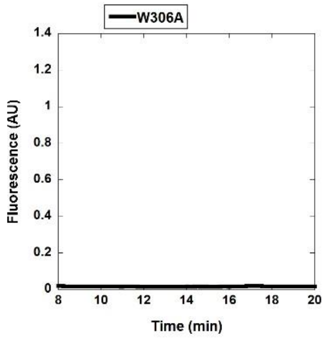
W306A
Fluorescence (AU)
Time (min)

n)

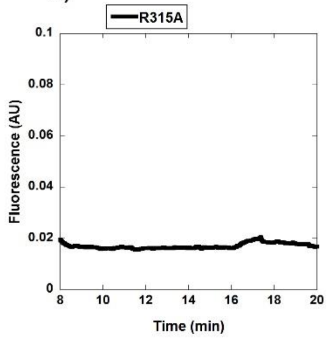
R315A
Fluorescence (AU)
Time (min)

o)

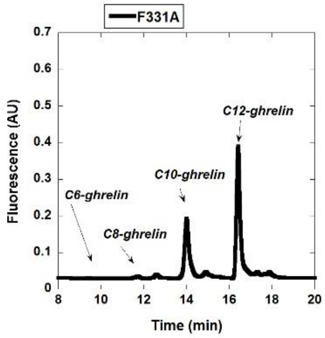
F331A
C6-ghrelin
C8-ghrelin
C10-ghrelin
C12-ghrelin
Fluorescence (AU)
Time (min)

p)

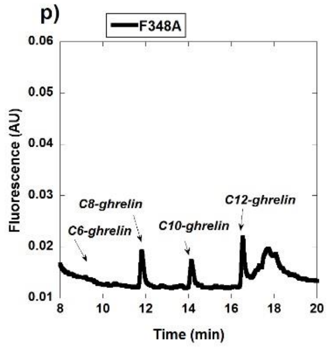
F348A
C6-ghrelin
C8-ghrelin
C10-ghrelin
C12-ghrelin
Fluorescence (AU)
Time (min)

q)

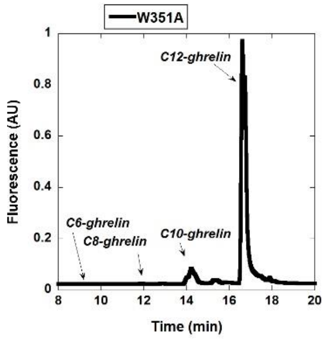
W351A
C6-ghrelin
C8-ghrelin
C10-ghrelin
C12-ghrelin
Fluorescence (AU)
Time (min)

r)

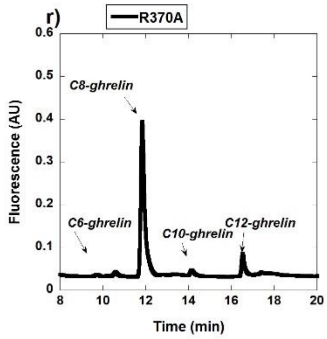
R370A
C6-ghrelin
C8-ghrelin
C10-ghrelin
C12-ghrelin
Fluorescence (AU)
Time (min)
